# ADAR mediated A-to-I RNA editing affects SARS-CoV-2 characteristics and fuels its evolution

**DOI:** 10.1101/2021.07.22.453345

**Authors:** Yulong Song, Xiuju He, Wenbing Yang, Tian Tang, Rui Zhang

**Affiliations:** MOE Key Laboratory of Gene Function and Regulation, Guangdong Province Key Laboratory of Pharmaceutical Functional Genes, State Key Laboratory of Biocontrol, School of Life Sciences, Sun Yat-Sen University, Guangzhou, 510275, PR China

## Abstract

Upon SARS-CoV-2 infection, viral intermediates activate the Type I interferon (IFN) response through MDA5-mediated sensing and accordingly induce ADAR1 p150 expression, which might lead to A-to-I RNA editing of SARS-CoV-2. Here, we developed an RNA virus-specific editing identification pipeline, surveyed 7622 RNA-seq data from diverse types of samples infected with SARS-CoV-2, and constructed an atlas of A-to-I RNA editing sites in SARS-CoV-2. We found that A-to-I editing was dynamically regulated, and on average, approximately 91 editing events were deposited at viral dsRNA intermediates per sample. Moreover, editing hotspots were observed, including recoding sites in the spike gene that affect viral infectivity and antigenicity. Finally, we provided evidence that RNA editing accelerated SARS-CoV-2 evolution in humans. Collectively, our data suggest that SARS-CoV-2 hijacks components of the host antiviral machinery to edit its genome and fuel its evolution.

## Main text

Severe acute respiratory syndrome coronavirus 2 (SARS-CoV-2), a positive-sense single-stranded RNA ((+)ssRNA) virus, emerged in late 2019 and expanded globally, resulting in over 82 million confirmed cases by the end of 2020 (*1, 2*). The ongoing evolution of SARS-CoV-2 has been a topic of considerable interest as the pandemic has unfolded, and new variants are continuing to spread (*3-6*). SARS-CoV-2 has complex processes of replication and transcription facilitated by the replication-transcription complex (RTC) with RNA-dependent RNA polymerase (RdRP) activity (*7*). Negative-strand RNAs are synthesized by RdRP starting from the 3’ end of positive genomic RNAs ((+)gRNAs), from which continuous synthesis generates full-length (-)gRNAs, whereas discontinuous jumping produces negative subgenomic RNAs ((-)sgRNAs). (+)gRNA and (+)sgRNA progenies are then synthesized using these negative-strand RNA intermediates as templates. During these processes, viral base substitutions are produced from two types of sources. The first type is substitutions introduced by replication error. SARS CoV-2 acquires such substitutions slowly as the result of a proofreading RdRp (*8*), and the mutation rate of replication error-introduced substitutions is estimated to be ∼3×10^−6^ per infection cycle (*9*). The second type is substitutions introduced by host-dependent factors, such as C-to-U and A-to-I RNA editing mediated by APOBEC and ADAR deaminases that target ssRNA and dsRNA substrates, respectively. As both ssRNA forms and dsRNA intermediates are present in the SARS-CoV-2 life cycle, both deaminases might bind and edit the SARS-CoV-2 genome.

In this study, we focused on ADAR-mediated A-to-I RNA editing (*10, 11*) and aimed to investigate the occurrence of A-to-I RNA editing and its impacts on SARS-CoV-2 characteristics and evolution. ADAR1 has a key role in suppressing IFN signaling and plays an important role during viral infections (*12-14*). Although signatures of ADAR activity have been found in a number of RNA viruses (*15*), no tool has been developed to systematically identify A-to-I RNA editing in viruses. Recent studies revealed that upon SARS-CoV-2 infection, viral intermediates activate the IFN response through MDA5-mediated sensing (*16*). The IFN response may further induce cytoplasmic expression of ADAR1 p150 isoform (*17*), which might lead to A-to-I RNA editing of SARS-CoV-2. A-to-I RNA editing in SARS-CoV-2 may regulate viral biology in two ways. First, viral editing within the infected individual may affect viral characteristics and virus-host interaction. In this case, we would expect that editing events need to achieve a certain level. e.g., at least a 5% level, for proper function. Second, viral editing as a possible source of mutations can fuel viral evolution during transmission. In this case, it is similar to the process in which newly emerged variants introduced by replication error are fixed at the point of transmission and promote the evolution of RNA viruses (*18, 19*). Even the sites with extremely low-level editing might contribute to viral evolution during viral spread.

Many tools have been recently developed to identify A-to-I RNA editing sites from RNA-seq data (summarized in (*20, 21*)), but detection typically requires matched genomic sequences from the same sample to discriminate RNA editing events from other types of variants. As RNA viruses do not possess an unedited DNA form, conventional pipelines, in principle, have limited power to detect editing sites in such samples. Indeed, an initial attempt (*22*) to identify A-to-I editing sites in SARS-CoV-2 using a conventional pipeline obtained putative variants with motifs inconsistent with the common view of ADAR mediated RNA editing, suggesting that most sites are false positives.

One hallmark of ADAR mediated editing is that editing sites are clustered together, and such a feature has been successfully used to develop a pipeline to call hyper-editing sites (*23*). Based on the cluster feature, we developed a computational pipeline that is specific for editing identification of RNA viruses (**Figure S1**, see details in **Methods**). The characteristic features that distinguish our approach are: (i) the careful design of read preprocessing and viral genome optimized mapping steps; (ii) the addition of a two-step editing calling step specifically for (+)ssRNA viruses, which may introduce both A-to-G and T-to-C variants in (+)gRNA and (+)sgRNA sequences during the production of (+)gRNAs and (+)sgRNAs (**Figure S2**); (iii) the incorporation of multiple additional filters to remove misaligned and low-quality reads; (iv) the editing site number filter to remove false-positives due to the overamplification issue of small-genome viruses; and (v) the editing level filter to remove variants that are unlikely introduced by an RNA editing mechanism.

To verify our pipeline, we applied it to RNA-seq data from a cell culture model of SARS-CoV-2 infection. We first confirmed that the conventional pipeline identified different types of RNA variants (**Figure S3A-B**), and the putative editing events (A-to-G variants and T-to-C variants (A-to-I RNA editing of the negative-strand RNA of SARS-CoV-2)) had no ADAR motif preference (**Figure S3C**). In sharp contrast, the vast majority of the variants identified with the RNA virus-specific pipeline were A-to-G/T-to-C types, indicative of A-to-I editing (**Figure S3D**). Most importantly, the nucleotides neighboring the A-to-G/T-to-C variants showed a pattern consistent with known ADAR preference (*24*), i.e., the underrepresentation of G upstream of the editing site (**Figure S3E**). Notably, the proportions of A-to-G/T-to-C sites and the motif preference were greatly increased with the edited read filtering steps of our pipeline (**Figure S3D-E**), supporting the efficacy of our filters. As expected, the proportion of viral reads and the number of editing sites increased with the increase of infection time (**Figure S3A & S3F**). Together, these data highlight the accuracy of our approach.

Having established the method for virus-specific editing calling, we analyzed RNA-seq data from diverse types of samples for a global characterization of A-to-I RNA editing events in SARS-CoV-2. A total of 7622 published RNA-seq samples, including nasopharyngeal swabs of COVID-19 patients, autopsy tissue samples of donors who died of COVID-19, and organoids and cell models infected with SARS-CoV-2, were collected (**Figure 1A & Table S1**). 1727 (23%) samples were found to have > 1000 reads mapped to the SARS-CoV-2 genome (**Figure 1B**), of which 227 samples had viral editing events (**Figure 1B**). On average, approximately 91 editing events were identified per sample. There was a clear positive correlation between the viral load and the detection of RNA editing (**Figure 1B**). For example, of the samples with viral reads > 5 million, 56% had editing events detected. The overall low percentage of edited reads could be explained by the possibility that A-to-I editing is effective in restricting viral propagation, thus reducing the number of viruses that show evidence of these changes. In total, we identified 8590 and 1415 A-to-G/T-to-C sites from the one-step and two-step editing calling pipelines (**Table S2**), respectively, and a substantial fraction of sites called from the two pipelines were overlapped (**Figure S4A**). As expected, editing sites from both pipelines showed a pattern consistent with known ADAR preference (**Figure 1C**). As a control, we called other types of variants using the same criteria (see **Methods**), and very few samples had other types of RNA variants detected (**Figure 1D**), further supporting the validity of our editing identification pipeline. Note that the use of the previously developed hyper-editing pipeline led to a large number of samples with other types of variants (e.g., A-to-T/T-to-A and A-to-C/T-to-G) identified (**Figure S4B**), highlighting the necessity of virus-specific filtering steps in our pipeline.

**Figure 1.**
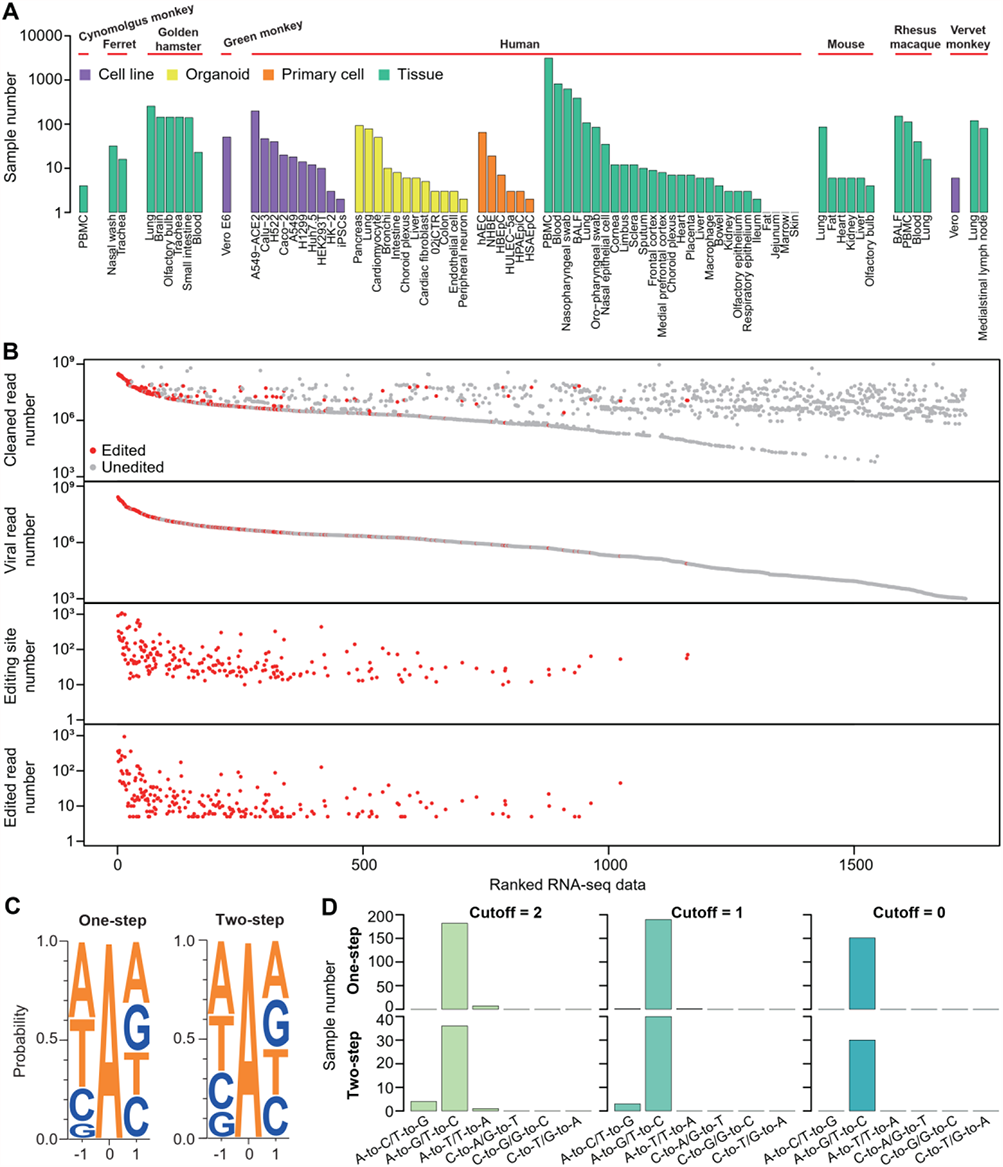
RNA virus-specific pipeline identifies authentic A-to-I RNA editing sites in SARS-CoV-2. **(A)** Summary of the RNA-seq data analyzed. PBMC, peripheral blood mononuclear cells; 02iCTR, a hiPSC line generated from peripheral blood mononuclear cells; hAEC, human aortic endothelial cells; NHBE, normal human bronchial epithelial cells; BALF, bronchoalveolar lavage fluid; HBEpC, primary human bronchial epithelial cells; HPAEpiC, human pulmonary alveolar epithelial cells; HSAEpC, primary human small airway epithelial cells. **(B)** The numbers of total cleaned reads, viral reads, editing sites, and edited reads in RNA-seq data that had viral reads > 1000. Samples were ranked by viral read number. **(C)** Nucleotides neighboring editing sites called using either one- or two-step editing pipelines showed a pattern consistent with known ADAR preference. The reverse complements of the triplets of T-to-C variants were combined with the triplets of A-to-G sites for analysis. The motif was characterized by the underrepresentation of G upstream of the editing site. **(D)** Numbers of samples with different variant types identified using our method with different mismatch and low-quality site cutoffs (see **Methods** and **Figure S1**).

Next, we investigated the landscape of A-to-I RNA editing in SARS-CoV-2. Similar numbers of A-to-G and T-to-C sites were identified, and both were relatively evenly distributed across the genome (**Figure 2A**). This result suggests that RNA editing mainly occurs in viral dsRNA intermediates, which can be produced during gRNA replication and sgRNA transcription processes (*7, 25*). In line with this, edited As did not tend to be located in the more structured regions as compared with the unedited As (**Figure 2B**). When examining the cluster features of editing sites, we found that most edited reads had fewer than six editing sites (**Figure 2C**), suggesting that extensively edited viruses may be defective and removed, consistent with the detection of overall low editing levels of SARS-CoV-2. There were a total of 3314 and 2018 A-to-G and T-to-C sites that can alter amino acids, which may potentially affect virus characteristics (**Table S2**). Note that some of the negative-strand RNAs with A-to-I editing (i.e. T-to-C variants) may not be further used as a template for (+)gRNA or (+)sgRNA synthesis and thus did not recode the viral genome.

**Figure 2.**
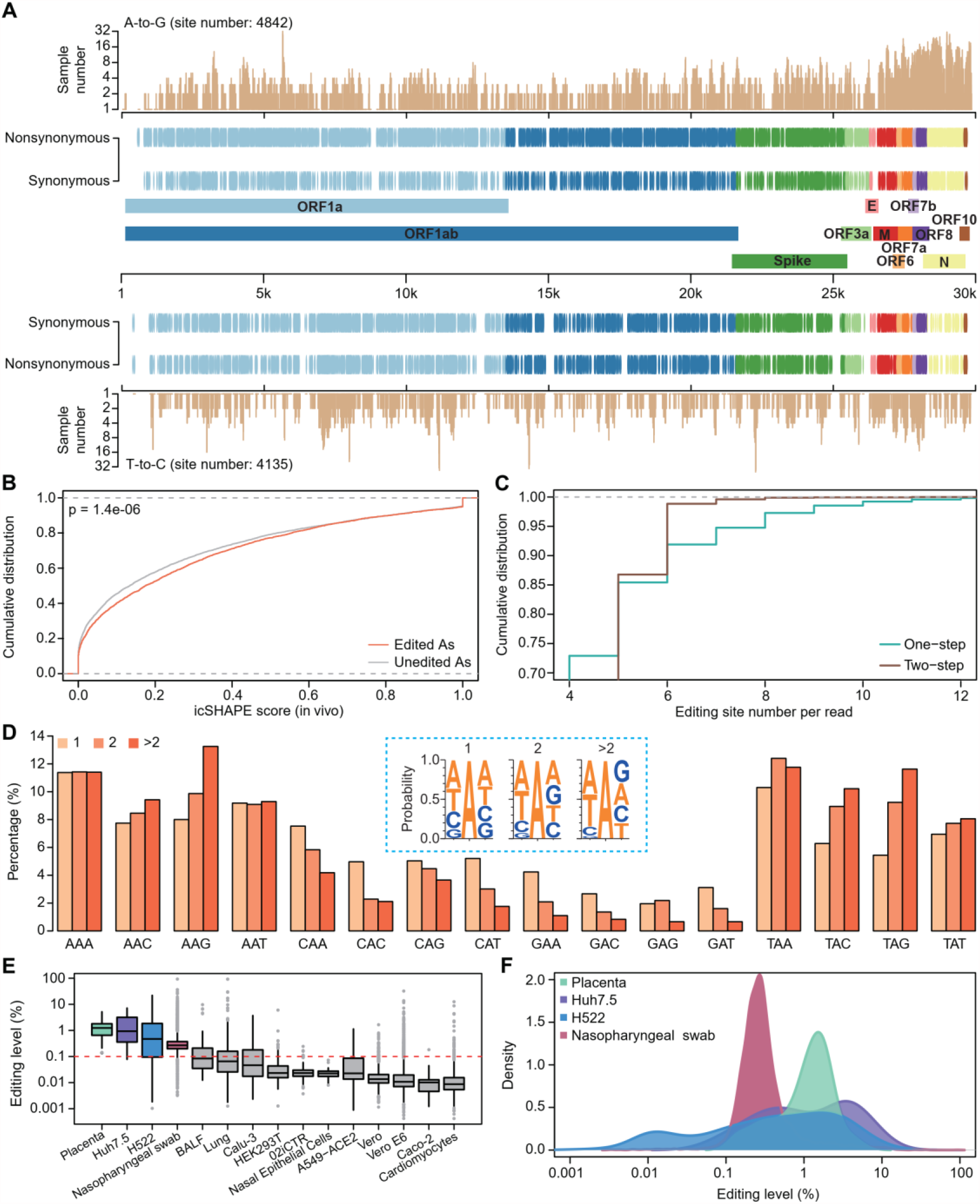
The landscape of A-to-I RNA editing events in the SARS-CoV-2 genome. **(A)** Genic location and annotation of editing sites SARS-CoV-2. A-to-G and T-to-C sites were plotted separately. The number of edited samples at each editing site is indicated. **(B)** Comparison of the icSHAPE scores between edited and unedited As in the (+)gRNA. A higher score indicates that a nucleotide is more likely single-stranded. P-value was calculated using Kolmogorov-Smirnov test. **(C)** The cumulative distribution of the number of editing sites per read. **(D)** Comparison of triplet preferences for editing sites identified in one, two, or more than two samples. The reverse complements of the triplets of T-to-C variants were combined with the triplets of A-to-G sites for analysis. **(E)** Boxplot showing the editing levels of sites among samples. Only editing sites with coverage ≥ 30 were used for analysis. Note that for the lowly edited sites (e.g. with editing level < 0.1%), our calculations overestimated the editing level because G reads caused by sequencing errors were also counted as edited reads (see **Methods**). Red line, sequencing error rate based on our quality cutoff (Q30). **(F)** The editing level distribution in selected human samples.

To ask whether editing hotspots were present in the viral genome, we examined the number of edited samples at each editing position. We found 2271 sites present in multiple samples, ranging from 3 to 41 (**Figure 2A**). As a group, the triplet motifs of editing hotspots were biased towards the ones most favored by ADAR1 (e.g., TAG) and biased against those unfavored by ADAR1 (e.g., GAN) (**Figure 2D**), which underlies the basis of their high editability.

To understand the dynamics of RNA editing, we performed three analyses. First, we compared the overall editing levels between samples and found that they varied (**Figure 2E**), which was probably due to the different levels of MDA5 associated response. Moreover, editing levels among sites varied within a sample (**Figure 2F**). Second, we examined the association between MDA5 expression and overall editing levels using cell models infected with SARS-CoV-2. A positive correlation between MDA5/ADAR1 expression and editing level was observed (**Figure S5**), suggesting that stronger innate responses may promote viral RNA editing. Third, we analyzed the viral editing status of 617 nasopharyngeal swab samples from the New York City metropolitan area during the COVID-19 outbreak in spring 2020 (*26*). We found that females had a higher viral editing index than males, particularly for those aged between 45 and 64 (**Figure S6**), which is consistent with previous observations that adult females mount stronger innate immune responses than males in general (*27*). Altogether, these data suggest that A-to-I RNA editing in SARS-CoV-2 was dynamically regulated.

The trimeric spike protein decorates the surface of coronavirus and plays a key role in viral infection and pathogenesis (*28*). It is also the major antigen inducing protective immune responses and the major target for vaccine development (*29, 30*). Thus, the editing-introduced mutations in the spike protein may provide a new source for its evolution and adaptation in humans. We found a total of 751 A-to-G/T-to-C recoding editing sites in the spike protein (**Figure 3A, Table S3**). A close inspection of these recoding sites revealed that these sites may affect virus characteristics in a variety of ways: 1) 151 sites were located at the linear epitope regions of the spike protein (*31*), which may affect the host immunogenic response (**Figure 3B**). 2) Although as expected that most recoding sites in the spike receptor binding domain (RBD) were deleterious for RBD expression and ACE2 binding, a few recoding sites were found to enhance RBD expression and ACE2 affinity (**Figure 3C**), based on deep mutational scanning data (*32*). 3) Four recoding sites were found to alter the ACE2 binding affinity to the spike protein and affect viral infectivity and antigenicity (**Figure 3D**), based on the high-throughput pseudovirus assay (*33*). 4) Computational prediction revealed that recoding sites tended to stabilize the protein (**Figure 3E**) and thus might have both pros and cons. On the one hand, recoding sites may enhance the spike-mAb interaction; on the other hand, they may also enhance the spike-ACE2 interaction. Altogether, RNA editing provides a new source to regulate virus-host interactions.

**Figure 3.**
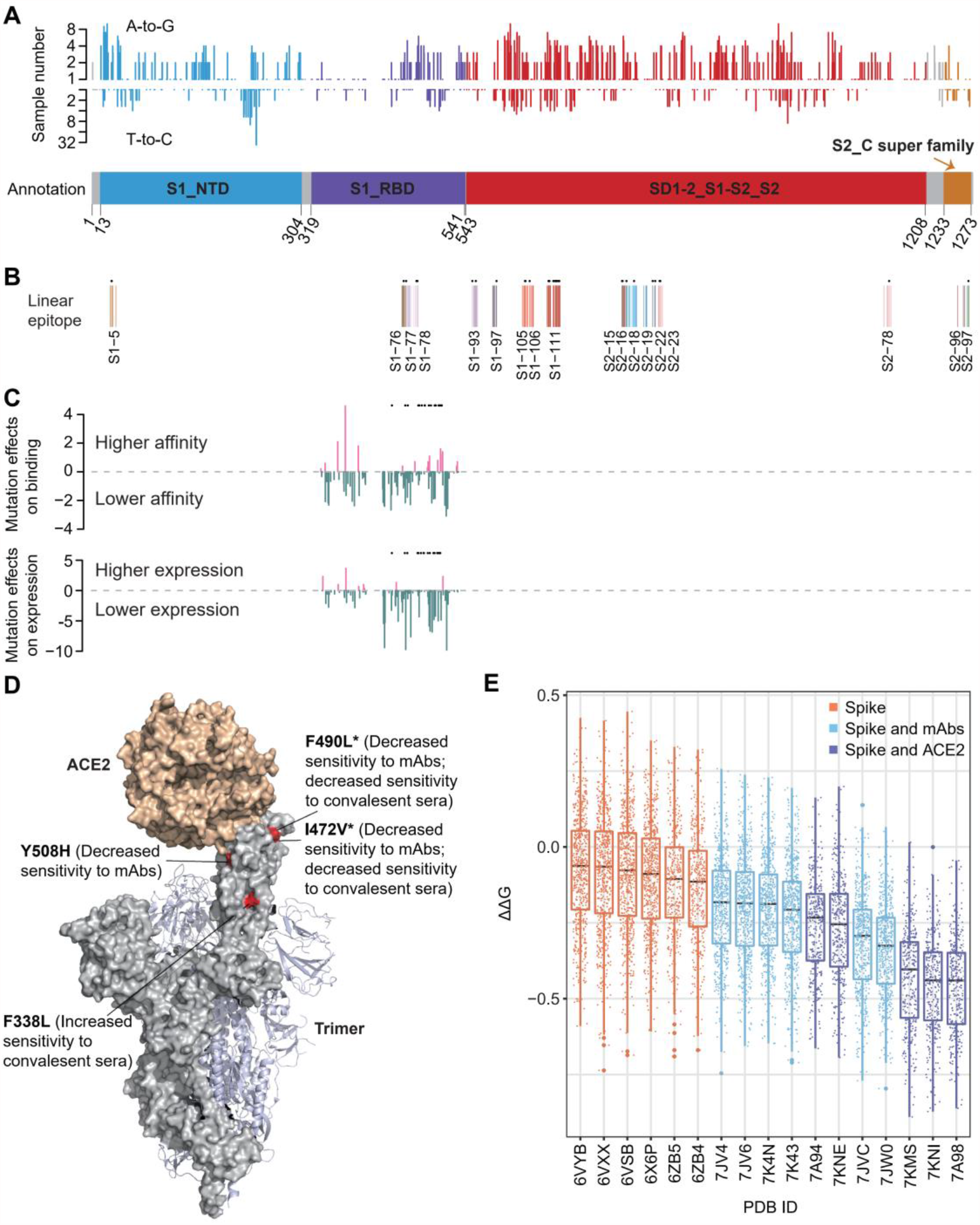
Impacts of RNA editing on spike protein functions. **(A)** The distribution of recoding editing sites in the spike protein. The number of edited samples at each editing site is indicated. **(B)** The recoding sites that were located at the linear epitope regions of the spike protein. Linear epitope regions were mapped by Li Y et al (*31*). The editing sites detected in more than two samples are labeled as “*”. **(C)** The effects of recoding editing on RBD expression and ACE2 binding. Data were from Starr et al (*32*). Positive value, higher affinity; negative value, lower affinity. The values were normalized to the median values of all positive or negative mutation effects. The editing sites detected in more than two samples are labeled as “*”. **(D)** Four recoding sites were found to affect viral infectivity and antigenicity. Data were from Li Q et al (*33*). Note that I472V recoding site in the D614G background can further lead to increased infectivity and decreased sensitivity to neutralizing mAb and convalescent sera. The editing sites detected in more than two samples are labeled as “*”. **(E)** The effects of recoding editing events on spike protein stability were predicted by MAESTRO based on the structures of the spike protein, spike-mAb complexes and spike-ACE2 complexes. Y-axis is the change in Gibbs free energy (ΔΔG) upon RNA editing. Positive and negative values represent the decrease and increase of the protein stability, respectively.

The RNA editing events may occur in both strands of sgRNAs and gRNAs. Those in sgRNAs may only affect the characteristics of the viruses within the infected individual; while those in gRNAs might transmit, thus fueling viral evolution in humans. Given the overall low-level editing in SARS-CoV-2, we expect that A-to-I editing may have a low or moderate effect on the infected individual. However, RNA editing introduced substitutions might provide a new source of mutation and contribute to SARS-CoV-2 evolution in human populations. Extensive global sampling and sequencing of SARS-CoV-2 have enabled us to examine the impact of A-to-I RNA editing on its evolution. Because A-to-I editing tended to be clustered, we reasoned that if A-to-I editing contributes to virus evolution, such cluster signature could be found in viral sequences. Thus, we searched the GISAID sequence database (*34*) for sequences containing clustered A-to-G mutations in the editing site positions (see **Methods**). As a control, we searched clustered A-to-C mutations in the same positions, because experimental data suggest that SARS-CoV-2 had comparable rates of replication error-introduced A-to-G and A-to-C mutations (*9*). We found 11-39-fold enrichment of sequences containing clustered A-to-G mutations for a cluster containing 2-6 mutations (**Figure 4A**), supporting our hypothesis. We also used randomly sampled unedited A positions as a control, and similar enrichment was observed (**Figure 4B**). If A-to-I RNA editing accelerates SARS-CoV-2 evolution in humans, we would expect that the edited A positions may be more likely fixed as Gs in the epidemic. We thus compared the frequencies of A-to-G mutations inferred using sequences in GISAID between edited and unedited As in the genome. We found that edited As tended to have higher frequencies, particularly for the sites in the intergenic regions (**Figure 4C**), which are expected to be under less selective constraints. Together, these analyses imply that RNA editing promotes SARS-CoV-2 evolution in humans.

**Figure 4.**
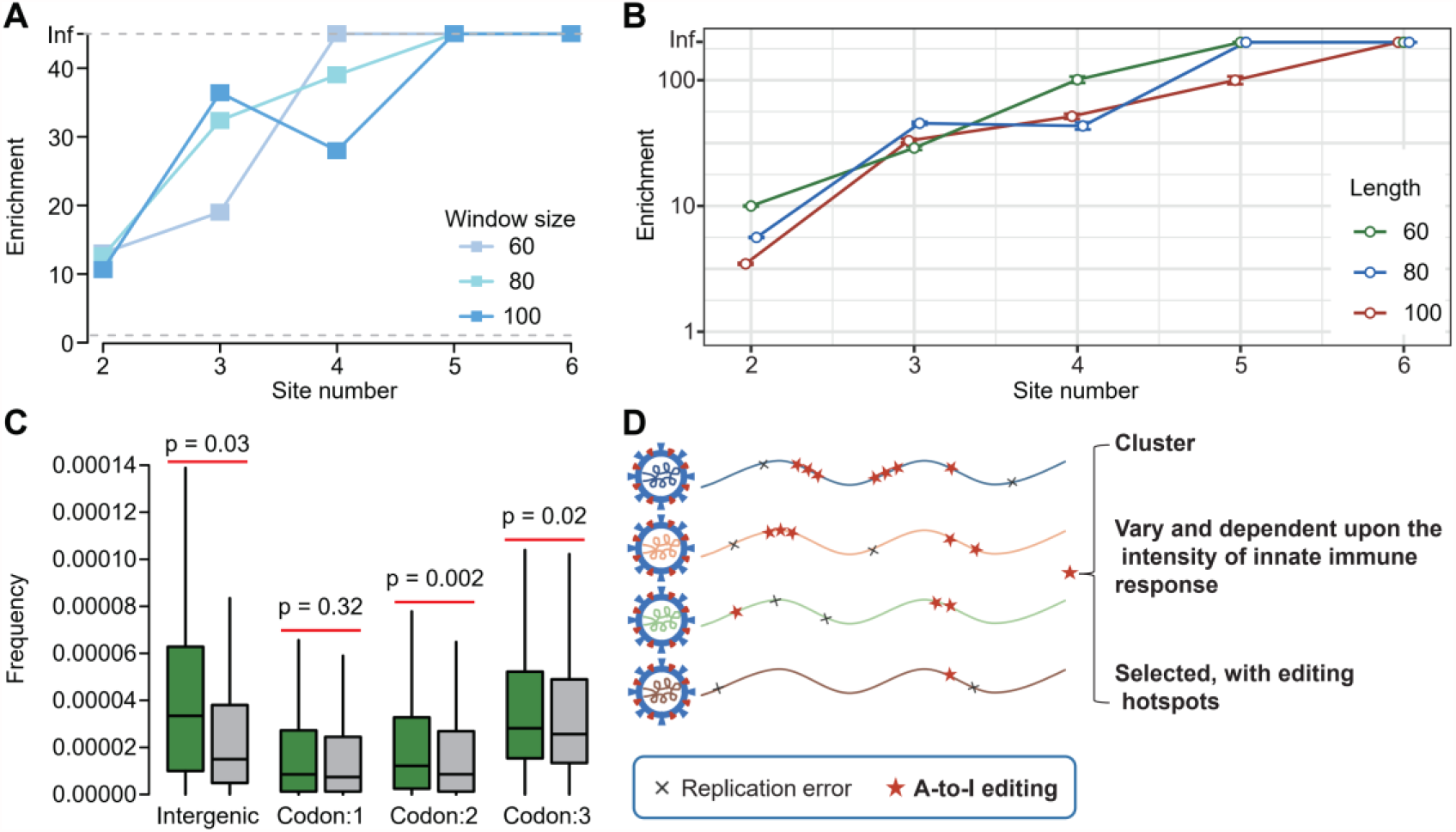
A-to-I RNA editing promotes SARS-CoV-2 evolution in humans. (**A-B**) Enrichment of the clustered A-to-G sites in the global samples of SARS-CoV-2. We used two different controls for enrichment calculation. In panel **A**, enrichment was defined as the number of clusters with A-to-G mutations in edited A positions divided by that with A-to-C mutations in edited A positions (see **Methods**). In panel **B**, enrichment was defined as the number of clusters with A-to-G mutations in edited A positions divided by that with A-to-G mutations in the randomly sampled unedited A positions (see **Methods**). Control sites were sampled 1000 times and 95% confidence interval is shown. In both **A** and **B**, “Inf” means no GISAID sequences in the control sets were identified. **(C)** Comparison of the GISAID A-to-G variant frequencies between edited and unedited As. Sites located in intergenic regions and three codon positions were analyzed separately. P-values were calculated using one-sided Mann-Whitney U-test. **(D)** Summary of the characteristic of A-to-I editing on SARS-CoV-2.

Knowledge of the host-dependent factors that introduce mutations is important for the simple reason that RNA viruses, particularly (+)ssRNA viruses, are major pathogens of humans, and we have had 3 coronavirus epidemics in the past 2 decades. In this study, we developed an approach to robustly identify A-to-I RNA editing in RNA viruses. Our approach paves the way for future studies of the impacts of A-to-I RNA editing on RNA virus biology. By applying our approach to SARS-CoV-2, we characterized the landscape and dynamics of RNA editing in the SARS-CoV-2 genome. We further revealed that A-to-I editing provides a new source of substitutions, and promotes the evolution of SARS-CoV-2 in humans. A comparison between our editing list and watch lists of variants of concern in the world revealed 20 overlapping sites (**Figure S7**), suggesting that such variants could be introduced by RNA editing.

Compared with replication error-introduced substitutions, A-to-I editing-introduced substitutions have distinct features (**Figure 4D**). First, RNA editing tends to be clustered, and such substitutions may lead to stronger changes in a small region, which provides a unique mode of substitutions for virus evolution. Second, editing levels vary among tissue types and between sexes, dependent upon the intensity of the innate immune response; thus, caution may be needed for individuals who have a stronger innate immune response and likely generate a high-level of editing-introduced A-to-G substitutions. Third, since ADAR has a motif preference, the editing sites are selected, and editing hotspots are presented. We propose that editing hotspots in the SARS-CoV-2 spike protein that enhance ACE2 affinity may be integrated into mRNA vaccine design in advance to defend against possible new variants.

## Methods

### The SARS-CoV-2 reference genome sequence

We used the complete genome sequence SARS-CoV-2 Wuhan-Hu-1 strain (Accession NC_045512, Version NC_045512.2) as the reference genome.

### RNA-seq data collection

RNA-seq data were downloaded from NIH SARS-CoV-2 resource website (https://www.ncbi.nlm.nih.gov/sars-cov-2/). A list of sample IDs is shown in **Table S1**. We only collected data generated using Illumina, Ion Torrent, and BGI platform.

### Mapping of RNA-seq reads

We first trimmed adapters and low-quality bases using cutadapt (*35*). Next, reads were trimmed to 60 - 80 nt, and reads with a length < 60 nt were discarded. The cleaned reads were mapped to the reference sequence using BWA (*36*) aln (bwa aln -t 20) and mem (bwa mem -M -t 20 -k 50). The unmapped reads were extracted for editing site calling.

### Calling of putative RNA editing sites

To realign reads with a cluster of mismatches caused by A-to-G editing, we transformed every A to G in both the unmapped reads and the viral reference genome. We aligned the transformed reads to the transformed genome, again using BWA aln (bwa aln -t 5 -n 2 -o 0). The original (four-letter) sequences of the reads that aligned (after the transformation) were recovered, and RNA variants between the reads and the reference genome were examined. For multiple mapped reads, only the best hits were retained. Next, edited reads and editing sites were called as described in **Figure S1**.

### RNA variant calling via the conventional editing identification pipeline

We used a widely used pipeline for conventional RNA variant calling (*37*). In brief, we mapped cleaned reads to the reference genome via BWA as above. Next, we removed identical reads (PCR duplicates). Then, we inspected all positions that showed variation in the RNA using samtools (Version: 1.2) mpileup. We only took variant positions in the RNA into consideration if they conformed to our requirements for the number and quality of bases that vary from the reference genome. We specifically required that each variant was supported by three or more variant bases having a base quality score ≥ 30. To avoid false positives at the 5’ read ends due to random-hexamer priming, we truncated the first 6 bases of each read. We also removed RNA editing candidates if they were located in regions of high similarity to other parts of the genome.

### Editing level quantification

We first merged the bam file containing clustered read mapping results with the bam file containing the mapped reads. Next, we inspected all editing positions using samtools for editing level quantification. We required that the minimal base quality was 30, and only sites covered by at least 30 reads were retained.

### Editing index analysis

617 nasopharyngeal swab samples from the New York City metropolitan area during the COVID-19 outbreak in spring 2020 (*26*) were used for analysis. For each sample, the editing index was defined as the number of edited viral reads divided by the total number of viral reads. Only reads with the number of mismatch and low-quality A-to-G/T-to-C sites ≤ 2 were considered as edited reads.

### Gene expression level quantification

TPM (Transcripts Per Million) was used to represent ADAR1 and MDA5 (*IFIH1* gene) expression. Reads were mapped to the human reference genome GRCh38 using Hisat2 (Version 2.0.4) (*38*) with default parameters. Then, StringTie (Version v1.3.0) (*39*) was used to calculate the TPM values with default parameters.

### Spike protein recoding site analysis

The effects of recoding sites in spike protein on RBD expression and ACE2 binding were measured based on deep mutational scanning data (*32*). The effects of recoding sites in spike protein on viral infectivity and antigenicity were measured based on the high-throughput pseudovirus assay (*33*). To evaluate the effects of RNA editing on spike protein, spike-ACE2, and spike-mAbs stability, all recoding sites in spike protein were used as the input mutation list, and impacts of single mutations on protein or protein complex stability were calculated by MAESTRO (*40*). PDB data (6VSB, 6VXX, 6VYB, 6X6P, 6ZB4,6ZB5, 7K43, 7K4N, 7A94,7A98, 7JV4, 7JV6, 7JVC, 7JW0, 7KMS, 7KNE, 7KNI (*41-45*)) were downloaded from Protein Data Bank.

To examine the overlaps between recoding editing sites in spike protein and variants of concern, we downloaded variant watch lists using the GISAID “Emerging Variants” portal.

### RNA structure analysis

The in vivo structure of SARS-CoV-2 was obtained from a previous study (*46*). For each nucleotide of the genome, an icSHAPE reactivity score between 0 and 1 for each nucleotide was obtained, with a higher score indicating that a nucleotide is more likely single-stranded.

### Protein structure illustration

The spike protein sequence diagram was created by IBS 1.0.3 (*47*). Spike protein crystallization data were from Zhou et al. (*45*) (PDB: 7KNE). Structure visualization and variant annotation were performed with PyMOL (http://www.pymol.org/).

### Motif discovery

For each RNA variant, the flanking sequence of each site was extracted from the reference genome. Motif logos were plotted with WebLogo v.3.5 (*48*).

### GISAID sequence analysis

Sequences were obtained from the GISAID database (*34*) and acknowledged in **Table S4**. Our dataset was composed of 827074 SARS-CoV-2 sequences collected and deposited between 12-1-2019 and 3-22-2021. All GISAID sequences were fist aligned to NC_045512.2 using blast (blastn -strand plus -num_threads 20 -line_length 300000 -perc_identity 80). The frequencies of A-to-G mutations were calculated based on the aligned data.

## Acknowledgments

We gratefully acknowledge the authors who generated and shared the raw RNA-seq data on which this research is based (**Table S1**). We also acknowledge the authors from the originating laboratories and the submitting laboratories, who generated and shared via GISAID genetic sequence data. We thank Dr. Chung-I Wu, whose speech on scientists should have a sense of social responsibility greatly encouraged the completion of this work. We also thank the colleagues in the microevolution WeChat group, particularly Dr. Chung-I Wu, Dr. Jian Lu, Dr. Xionglei He, Dr. Xiaowei Jiang and Dr. Haipeng Li, whose discussions on SARS-CoV-2 spread and evolution inspired this study.

## Funding

This study was supported by grants from Guangdong Major Science and Technology Projects (2017B020226002 to R.Z.), Guangdong Innovative and Entrepreneurial Research Team Program (2016ZT06S638 to R.Z.), National Natural Science Foundation of China (31900437 to Y.L.S.), Guangdong Science and Technology Department (2020B1212060031 to R.Z.; 2021A1515012463 to W.B.Y.) and Chinese Postdoctoral Science Foundation (2020TQ0389 to W.B.Y.).

## Author contributions

R.Z. designed the experiments; Y.L.S., X.J.H., W.B.Y. analyzed data; R.Z. and Y.L.S. wrote the manuscript with input from all authors.

## Competing interests

The authors declare no competing financial interests.

## Notes

### Competing Interest Statement

The authors have declared no competing interest.

